# A Yersinia effector activates JAK-STAT signaling in human macrophages

**DOI:** 10.1101/2023.05.08.539843

**Authors:** Laura Berneking, Indra Bekere, Marie Schnapp, Jiabin Huang, Klaus Ruckdeschel, Martin Aepfelbacher

**Affiliations:** Institute of Medical Microbiology, Virology and Hygiene, University Medical Center Hamburg-Eppendorf (UKE), Hamburg, Germany

**Author notes:** Corresponding author: Institute of Medical Microbiology, Virology and Hygiene, University Medical Center Hamburg-Eppendorf (UKE), Martinistr. 52, D-20246 Hamburg, Germany.

**Keywords:** Yersinia, JAK-STAT signaling, Stat, IL-10 signaling

## Abstract

The multifunctional *Yersinia* effector YopM inhibits effector triggered immunity and increases production of the anti-inflammatory cytokine Interleukin-10 (IL-10) to suppress the host immune response. Previously it was shown that YopM induces IL-10 gene expression by elevating phosphorylation of the serine-threonine kinase RSK1 in the nucleus of human macrophages. Using transcriptomics, we now show that YopM affects expression of genes encoding components of the JAK-STAT signaling pathway. Further analysis revealed that YopM mediates nuclear translocation of the transcription factor Stat3 in *Y. enterocolitica* infected macrophages and that knockdown of Stat3 inhibited YopM-induced IL-10 gene expression. YopM-induced Stat3 translocation did not depend on autocrine IL-10, activation of RSK1 or tyrosine phosphorylation of Stat3. Thus, besides activation of RSK1, stimulation of nuclear translocation of Stat3 is another mechanism by which YopM increases IL-10 gene expression in macrophages.

## Introduction

To subdue the immune response of the host for their own purposes, a variety of pathogens modulate the production of pro-inflammatory, but also anti-inflammatory cytokines by innate immune cells (Mogensen 2009, Asrat et al. 2015). For this purpose, various gram-negative bacteria including *Pseudomonas*, *Salmonella*, *Shigella* and *Yersinia* employ a molecular injection machine (termed type III secretion system / injectisome) to introduce pathogen specific effector proteins into neutrophils, macrophages and dendritic cells (Marketon et al. 2005, Finlay et al. 2006, Galan et al. 2006). The bacterial effector proteins often have unique molecular properties through which they can positively or negatively regulate key signaling pathways in host cells (Aktories et al. 2005).

Human pathogenic *Yersinia* spp. (*Y. enterocolitica, Y. pseudotuberculosis* and *Y. pestis*) use up to seven effector proteins (YopH, YopO, YopP/J, YopE, YopM, YopT and YopQ/K) two of which, YopP/J and YopM directly interfere with cytokine expression and -production by innate immune cells (Aepfelbacher et al. 2007, Dewoody et al. 2013). YopP/J is an acetyltransferase that acetylates and inhibits components of NF-κB and MAPK pathways, very efficiently inhibiting cytokine gene expression and -production (Ruckdeschel et al. 1997, Orth et al. 1999, Mukherjee et al. 2006, Schubert et al. 2020). YopM is a leucine-rich repeat scaffold protein which has been reported to inhibit the production of the proinflammatory cytokines Interleukin 1β (IL-1β) and tumor necrosis factor-α (TNF-α) in macrophages and mice (Kerschen et al. 2004, McPhee et al. 2010, LaRock et al. 2012, Chung et al. 2014, Hofling et al. 2014). Interestingly, in contrast to YopJ, YopM increases serum levels of the immunosuppressive cytokine Interleukin-10 (IL-10) in *Yersinia* infected mice, and this was found to be important for the virulence of the bacteria (Auerbuch et al. 2007, McPhee et al. 2010, McPhee et al. 2012). The IL-10-inducing effect of YopM in mice depends on its interaction with the host cell kinases RSK1 and PRK2 (McPhee et al. 2010, McPhee et al. 2012). Recently a molecular mechanism putatively underlying the IL-10-inducing effect of YopM was described. YopM was shown to stimulate phosphorylation of ribosomal S6 kinase 1 (RSK1) in the nucleus of human macrophages and this correlates with IL-10 gene expression (Berneking et al. 2016).

IL-10 is a key immunosuppressive cytokine produced by numerous immune cells, including macrophages. It mainly acts by downregulating the expression of inflammatory cytokines (Fiorentino et al. 1989, de Waal Malefyt et al. 1992, Chang et al. 2000, Staples et al. 2007, Saraiva et al. 2010). In this way, IL-10 plays a central role in regulating the balance between pro and anti-inflammatory responses in infections, which is also required for mounting an effective immune response (Mosser et al. 2008). Macrophages, monocytes, and dendritic cells all produce IL-10 in response to microbial pathogens, following their detection by pattern recognition receptors (PRRs), such as toll like receptors (TLRs). IL-10 is produced during infection with a number of bacterial, parasitic and viral pathogens, e.g., *Mycobacterium tuberculosis*, *Escherichia coli*, *Streptococcus pyogenes*, *Leishmania major*, and Cytomegalovirus, has in special circumstances been shown to support the virulence strategy of the pathogens, probably because it suppresses the antimicrobial effector functions of proinflammatory cytokines (Carey et al. 2012).

The effect of IL-10 on immune cell functions is exerted through its binding to and activation of the IL-10 receptor complex (IL-10R1 and IL-10R2), which stimulates Janus associated kinase (JAK)-STAT signaling (Riley et al. 1999). JAKs recruited to the activated IL-10 receptor tyrosine phosphorylate STAT transcription factors, such as Stat1 and Stat3, which thereby dimerize and translocate to the nucleus (Imada et al. 2000, Murray 2005). Stat3 that specifically transmits the signal of the activated IL-10 receptor stimulates transcription of various genes whose products exert anti-inflammatory activities (Mosser et al. 2008). Stat3-induced genes include the suppressors of cytokine signaling 1 and 3 (Socs1 and Socs 3) which effectively inhibit inflammatory signal transduction, including JAK-STAT signaling itself (excluding the IL-10R1/R2 triggered JAK-STAT signaling) and NF-κB and MAPK pathways (Imada et al. 2000). Another Stat3-induced gene is IL-10 itself, as part of a positive feedback loop (Staples et al. 2007). The activities of STAT3-induced genes together result in reduced production of proinflammatory mediators like TNF-α, IFN-γ, IL-6, and nitric oxide during an infection (Johnston et al. 2003, Yang et al. 2007).

Interestingly, some pathogens do not appear to directly induce and/or utilize IL-10 but have found ways to activate STAT3 in the IL-10 signaling pathway as part of their virulence strategy (Duell et al. 2012). The intracellular parasite *Toxoplasma gondii* has been shown to activate Stat3 through its effector rhoptry kinase 16 (Butcher et al. 2011). Further, the Salmonella effector SarA/SteE was found to phosphorylate and activate Stat3 by mimicking the cytoplasmic domain of glycoprotein 130 (gp130, IL6ST), a cytokine receptor that physiologically stimulates JAK-STAT signaling. *Salmonella* SarA is tyrosine phosphorylated in host cells, inducing the formation of a complex of Stat3 and GSK-3, whereby the latter, but not JAKs, mediate the phosphorylation and activation of Stat3 (Gibbs et al. 2020). In the last years it has been worked out in detail that expression of the IL-10 gene can be induced in different cell types by a multitude of signaling pathways, e.g., JAK-STAT, NF-κB, ERK-, p38 MAP and RSK kinases, which activate numerous transcription factors (Saraiva et al. 2010). These include e.g., Stat3 as well as CREB and ATF known to be activated by RSK-kinases downstream of ERKs (Anjum et al. 2008, Saraiva et al. 2010, Cargnello et al. 2011). In this work we searched for further mechanisms of *Yersinia*-induced IL-10 gene expression. We found that YopM triggers nuclear translocation of the JAK-STAT transcription factor Stat3 in human macrophages. This identifies a novel molecular mechanism by which YopM stimulates IL-10 gene expression.

## Results

### T3SS effector YopM maintains an elevated level of IL-10 in *Y. enterocolitica* infected macrophages through activation of RSK kinase

In previous work we described a molecular mechanism whereby YopM regulates the phosphorylation of Ribosomal S6 Kinase 1 (RSK1) in the nucleus of *Y. enterocolitica* infected human macrophages to control IL-10 gene expression (Berneking et al. 2016). Here, we aimed to elaborate on this mechanism and looked for possibly additional and/or synergistic mechanisms by which multifunctional YopM induces IL-10 in macrophages. For this, we infected macrophages with mock (uninfected), the avirulent strain WAC, wildtype strain WA314 and strains lacking YopP (WA314ΔYopP) or YopM (WA314ΔYopM) for 1.5 h and 6 h followed by RNA-Seq and selected qPCRs (Methods; S1 Table Fig. 1A). The avirulent strain WAC, known to trigger a strong PAMP-induced inflammatory cytokine production in macrophages, caused a 20-fold and 45-fold upregulation of IL-10 gene expression (i.e., IL-10 RNA levels) vs. mock after 1.5 h and 6 h of infection, respectively (Fig. 1B). It has been reported that in LPS-stimulated macrophages induction of the anti-inflammatory cytokine IL 10 occurs as a counter-response to excessive production of inflammatory cytokines and inhibits their activities (Carey et al. 2012). Wild type strain WA314, which strongly dampens the macrophage inflammatory response, showed only a minor IL-10 induction, amounting to about 2-fold the value of mock at 6 h (Fig. 1B). In turn, the WA314ΔYopP strain, which lacks the major immunosuppressive effector YopP, significantly induced expression of IL-10 again (approximately 10-fold the value of mock at 6 h (Fig. 1C). Notably, the WA314ΔYopM strain induced lower IL-10 mRNA values than wild type WA314 and showed no IL-10 increase compared to mock (Fig. 1B). This shows that the residual increased IL-10 mRNA expression in cells infected with wild type is caused by YopM. To test whether the IL-10 mRNA expression changes are reflected by IL-10 protein changes, we performed an IL-10 ELISA. The avirulent WAC induced high levels of IL-10 protein, which were reduced by 90% by wild-type WA314, with YopP playing the major role (Fig. 1C). Again, YopM was responsible for much of the IL 10 protein upregulated in macrophages infected with wild type bacteria (Fig. 1C). We conclude that along with the strong inhibition of inflammatory gene expression, also the expression of anti-inflammatory IL-10 is abolished in wild type *Y. enterocolitica* infected macrophages. However, it appears that YopM thereby counteracts the otherwise complete inhibition of IL-10 production by the *Yersinia* effectors. Although the YopM-induced upregulation of IL-10 mRNA expression was only two-fold and thus appeared relatively small compared to the strong IL-10 mRNA induction by WAC infection (1B), it resulted in an approximately doubled amount of IL-10 protein produced by the macrophages (Fig. 1C). Elevated levels of IL-10 appear to be beneficial for the propagation of *Yersinia* infection (Sing et al. 2002, McPhee et al. 2010, McPhee et al. 2012), a phenomenon also seen in other bacterial infections (van Der Poll et al. 1997, Duell et al. 2012).

**Figure 1.**
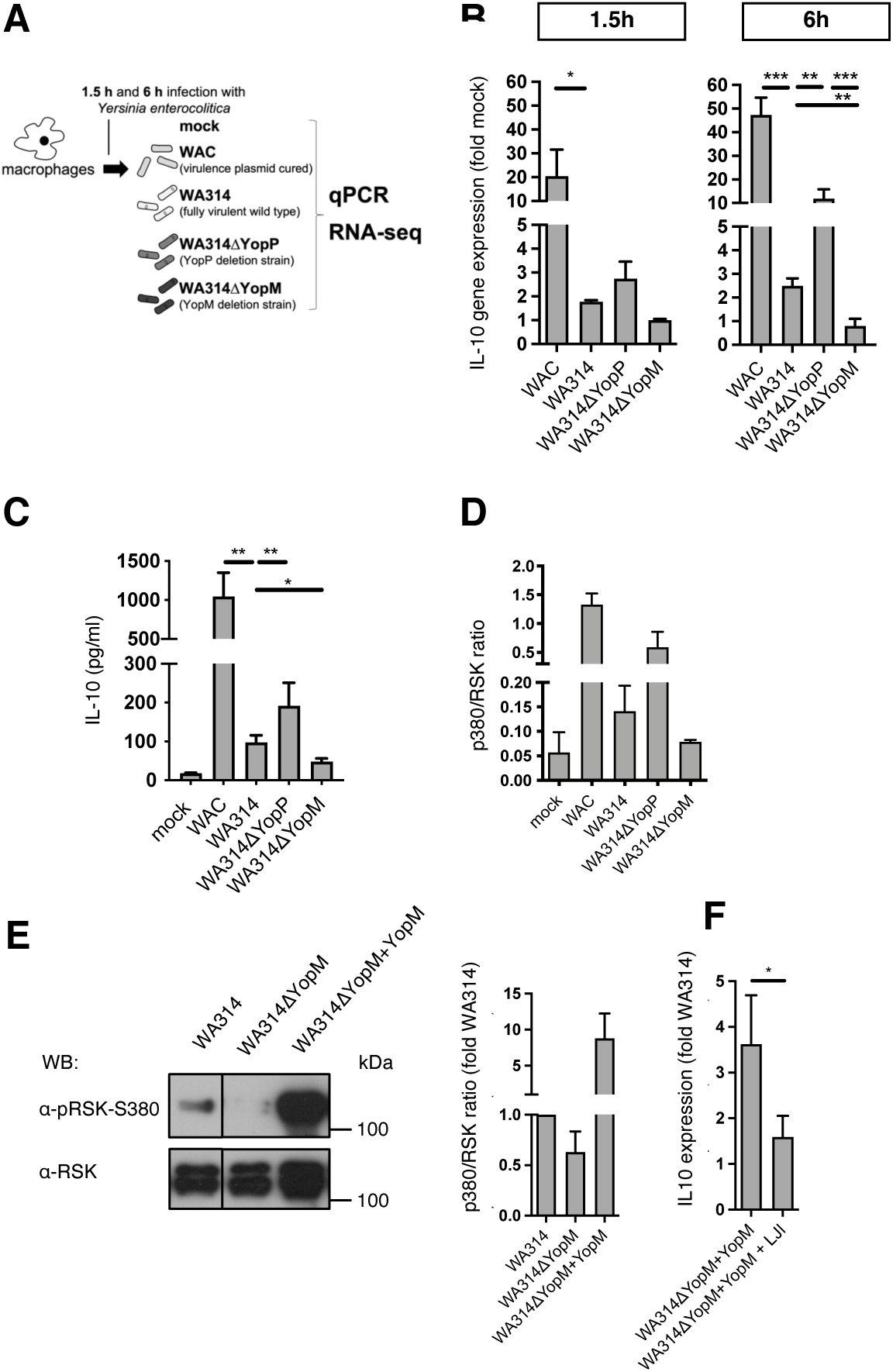
YopM induces the expression and production of IL-10. **A)** Experimental design. Monocytes were differentiated to human macrophages in the presence of human serum and infected after 7 days for 1.5 or 6 hours with *Yersinia* strains WAC, wildtype WA314, WA314ΔYopM, WA314ΔYopP or treated with PBS (mock). **B)** Total RNA was isolated from primary human macrophages infected for 1.5 or 6 h with indicated *Yersinia* strains WAC, WA314, WA314ΔYopM, WA314ΔYopP and subjected to RT-qPCR using human IL-10 specific primers. Expression was normalized to expression of three different housekeeping genes (GAPDH, TBP, B2M). For each condition triplicate samples of macrophages derived from different donors were investigated. Each bar in graph represents mean ± SEM of values from all donors normalized to mock. Analysis was performed with a linear mixed model considering random intercept. Data was transformed to ln-Score to ensure normal distribution; *p<0.05, **p<0.01, ***p<0.001. **C)** Human macrophage samples from four donors were infected with different Yersinia strains and supernatants were harvested at 6 h post infection for analysis of IL-10 by ELISA. Results shown are the mean of four independent experiments with error bars representing standard deviation. **D-E)** Human macrophages were infected with indicated *Yersinia* strains for 1.5h. Whole cell lysates were investigated by immunoblot using indicated antibodies. The phosphorylation of amino acid S380 of RSK1 was analyzed and compared to levels of RSK1. Intensities of the pS380 and RSK1 protein bands were quantified to calculate the intensity ratios of pS380/RSK1. **F)** Analysis of IL-10 gene expression in *Yersinia* infected human macrophages during LJI treatment. Primary human macrophages were treated with DMSO (Sigma) or RSK inhibitor LJI308 (LJI; 10µM, Sigma) for 1h and subsequently infected with WA314, WA314ΔYopM+YopM for 6 h. Total RNA was isolated and subjected to quantitative RT qPCR using human IL-10 specific primers. Expression of IL-10 was normalized to housekeeping genes GAPDH, TBP and B2M. For each condition triplicate samples of macrophages derived from three different donors were investigated. Each bar represents mean ± SEM of values from all donors.

In a previous study, we showed that a YopM-dependent increase of Ser-380 phosphorylation (p380) in RSK1 was associated with stimulation of IL-10 mRNA expression (Berneking et al. 2016). When we compared the levels of p380 RSK1 and IL-10 production in the macrophages infected with the different *Yersinia* strains here, we confirmed that they correlated closely (compare Fig. 1C and 1D). Because the differences in p380 RSK1 between WA314 and WA314ΔYopM were relatively small (Fig. 1D), we employed strain WA314ΔYopM+YopM, that translocates 2-3 times higher amounts of YopM into target cells than wild type (Hentschke et al. 2010, Berneking et al. 2016). Compared to WA314, WA314ΔYopM+YopM induced an approximately 9-fold increase of p380 RSK1, which was accompanied by a more than 3-fold increase in IL-10 mRNA levels (Fig. 1E and F). Using this enhanced effect of WA314ΔYopM+YopM on IL-10 mRNA expression, we could experimentally demonstrate that the pan-RSK inhibitor LJI308 significantly inhibits YopM-triggered IL-10 mRNA expression (Aronchik et al. 2014, Davies et al. 2015) (Fig. 1F). That LJI308 principally inhibits RSK kinases in primary human macrophages was demonstrated by inhibition of WAC-induced Histone-3-Ser-10 phosphorylation (S Fig. 1A), which is an RSK-dependent process triggered by WAC/LPS signaling (Sassone-Corsi et al. 1999); (S Fig. 1A). With this experiment we show a functional relationship between *Yersinia*-induced p380 RSK phosphorylation and IL-10 gene expression.

Overall, we show that YopP suppresses PAMP-induced IL-10 gene expression in *Y. enterocolitica*-infected macrophages and that YopM counteracts this YopP effect, to which upregulation of RSK kinase phosphorylation contributes.

### YopM modulates JAK-STAT signaling in macrophages

To gain more insights into the mechanisms of YopM-triggered IL-10 induction, we investigated the contribution of YopM to the transcriptional response of *Yersinia* infected macrophages. For this we focused our analysis on genes that were previously shown to be upregulated by IL-10 signaling in human monocytes (referred to as IL-10-dependent; (Teles et al. 2013). Correlating with the extent of IL-10 induction, 189, 108, and 40 IL-10-dependent genes were upregulated by WAC, WA314, and WA314ΔYopM, respectively (Fig. 2A, S2 Table). The number of IL-10-dependent genes upregulated by the different strains corresponded to approximately 5% of total upregulated genes (exact numbers in Methods). When the IL-10-dependent upregulated genes were subjected to KEGG pathway analysis, JAK-STAT signaling, FoxO signaling, cancer pathways and osteoclast differentiation were identified as enriched (Fig. 2B). Interestingly, the JAK-STAT and cancer pathways enriched in the WA314 upregulated genes showed lower p-values in the WA314ΔYopM upregulated genes (Fig. 2B; S3 Table), suggesting that the JAK-STAT pathway may be specifically induced by YopM. In fact, specific genes upregulated by WA314 but not by WA314ΔYopM include *IL10*, *JAK3*, *PTPN2* and *SOCS3* (list of all genes in S2 Table) all belonging to JAK-STAT pathway (Fig. 2B, S3 Table). A Venn diagram depicting the numbers of IL-10-dependent genes upregulated in WAC-, WA314- and WA314ΔYopM infected macrophages showed that altogether 78 genes (67+11 in green circle) were upregulated by WA314 but not by WA314ΔYopM, indicating that they were specifically induced by YopM (Fig. 2C; S4 Table). Pathway analysis of these genes again included JAK-STAT signaling (Fig. 2C; S5 Table).

**Figure 2.**
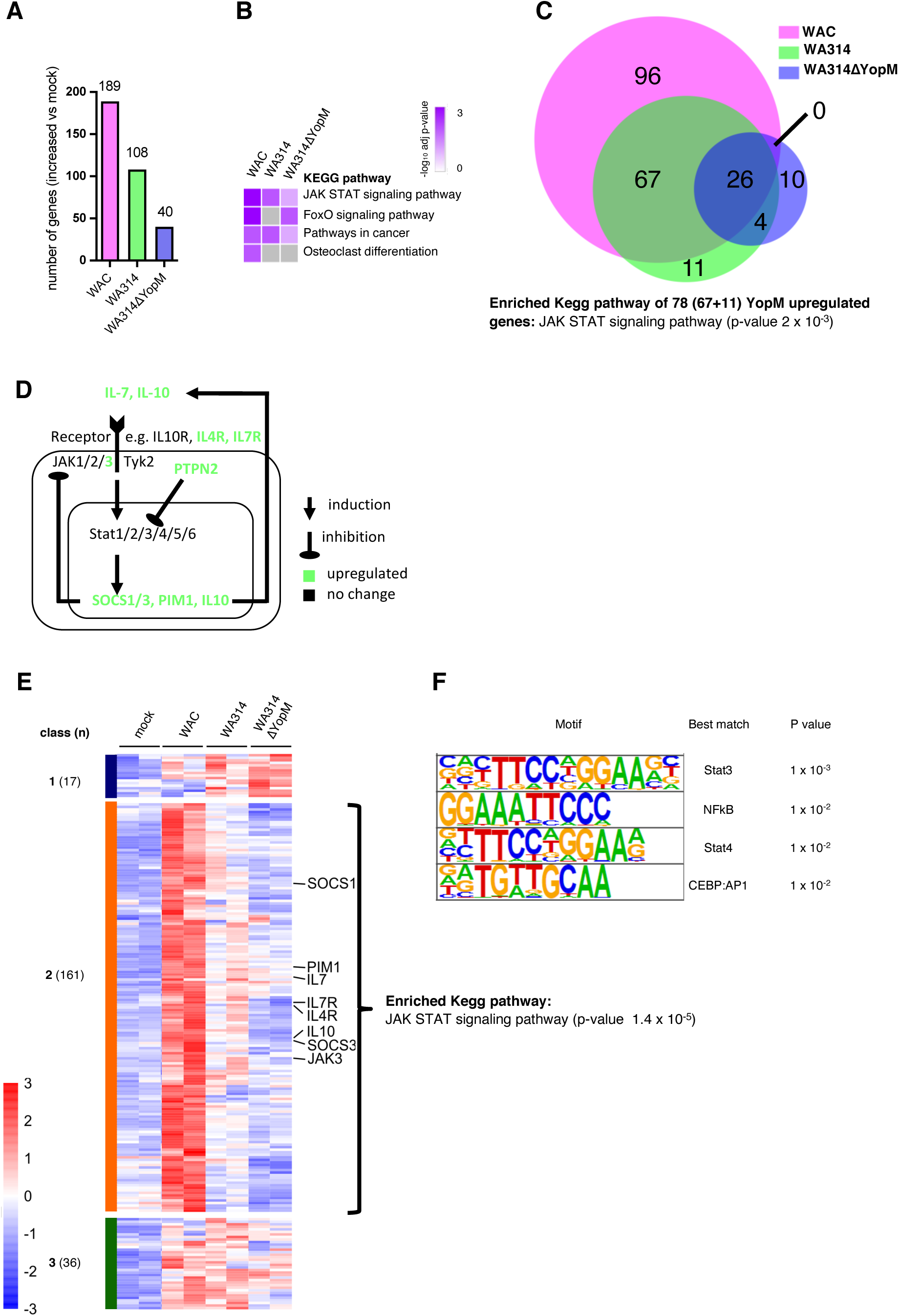
IL-10 induced transcriptional regulation by TTSS effector YopM. **A)** Differentially expressed genes (DEGs; log2fold change>1; adjusted p-value>0.05) in indicated infection-condition compared to mock that are IL-10-dependent as obtained from GSE43700 (Teles et al. 2013). **B)** Heatmap showing the *P*-value significance of distinct KEGG pathway enrichment for genes in each condition in (A). **C)** Venn Diagram of DEGs from mock, WAC, WA314 and WA314ΔYopM; the sum of the numbers in each circle represents total number of DEGs between combinations; the overlapping part of the circles represents common DEGs. **D)** Modulation of the JAK-STAT signaling pathway by YopM during *Yersinia* infection, DEGs are depicted in green. **E)** Heatmap from clustering of all IL-10-dependent DEGs from mock, WAC, WA314 and WA314ΔYopM comparisons at 6 h. Clustering identified 3 major classes. Rlog counts of DEGs were row-scaled (row Z-score). **F)** TF motif enrichment associated with DEGs in (E).

The JAK-STAT signaling cascade is initiated by binding of IL-10 and numerous other ligands to their cell surface receptors which recruit intracellular Janus kinases (JAK1, 2, 3) leading to phosphorylation, dimerization and finally nuclear translocation of STAT transcription factors ((Imada et al. 2000), Fig. 2D). One of these transcription factors, Stat3, is phosphorylated at Tyr-705 triggering its translocation into the nucleus. pTyr-705 Stat3 induces transcription of IL-10 as part of a positive feedback loop and Suppressor of Cytokine Signaling genes (SOCS), as part of negative feedback loops in JAK-STAT signaling (Berlato et al. 2002, Williams et al. 2004, Murray 2007, Jenkins 2014). Protein tyrosine phosphatases (PTPs) are further negative regulators of JAK-STAT signaling (Seif et al. 2017) whereby PTPN2 specifically reduces the phosphorylation level of Stat3 at Tyrosine 705 (Y705) resulting in decreased expression of pro inflammatory cytokines and cell proliferation (Zhang et al. 2018, Svensson et al. 2019).

In order to analyze the relative expression changes of IL-10-dependent genes in more detail, we created a heatmap of all respective DEGs in mock-, WAC-, WA314- and WA314ΔYopM infected macrophages. Clustering analysis revealed altogether 214 DEGs in three classes, characterized by distinct patterns of gene expression (classes 1, 2 and 3, Fig. 2E; S6 Table). Class 1 genes (n=17, blue color code) were upregulated by WA314 and more so by WA314ΔYopM, indicating that YopM downregulates expression of these genes (Fig. 2E). Class 3 genes (n=36, green color code) were upregulated to about the same extent by WAC, WA314 and WA314ΔYopM, indicating that they are regulated independently of YopM (Fig 2D). Of note, class 2 genes (n=161, orange color code) were upregulated by WAC and this was reduced by WA314 and more so by WA314ΔYopM (Fig. 2E), indicating that these genes are downregulated by WA314 effectors but upregulated by YopM. In class 1 and 3 genes no pathways were specifically enriched. Class 2 genes were enriched in JAK-STAT signaling pathway (Fig. 2E; genes belonging to JAK-STAT signaling are highlighted at the right and in green letters in Fig. 2D, S7 Table). Consequently, analysis of transcription factor motif enrichment of class 2 genes revealed binding sites for STAT- (Stat3/4), RHD- (NFkB-p65-Rel) and bZIP- (C/EBP, AP-1) families (Fig. 2F). Collectively, these data indicate that YopM induced upregulation of IL-10 is associated with the upregulation of numerous IL-10-dependent genes, among which components of the JAK-STAT pathway are highly enriched. Because the JAK-STAT pathway itself also induces IL-10 gene expression, these data raised the possibility that JAK-STAT signaling directly contributes to YopM-induced IL-10 induction.

### YopM promotes nuclear translocation of Stat3

To further analyze in which way JAK-STAT signaling may contribute to YopM-triggered IL 10 induction, we assayed translocation of Stat3 and Stat1 to the nucleus in macrophages infected with WAC, WA314 or WA314ΔYopM for 1.5 h and 3 h (Fig. 3A; S Fig. 2A and S Fig. 3). The degree of nuclear localization of Stat3 was quantified using immunofluorescence staining and Western blot of cell fractions (Fig. 3A-C). By immunostaining, Stat3 was found almost exclusively in the cytoplasm of uninfected cells (mock) (about 7 % Stat3-positive nuclei) and it partly shifted to the nucleus upon WAC infection (about 32 % Stat3-positive nuclei; Fig. 3A and B). Nuclear Stat3 increased to even higher values in WA314 infected macrophages, (about 55 % Stat3-positive nuclei), but in WA314ΔYopM infected cells it was strongly reduced, corresponding to the levels of uninfected cells (about 9 % Stat3-positive nuclei; Fig. 3A and B). This observed pattern of nuclear localization of Stat3 was similar at 1.5 h and 3 h of infection (S Fig. 2A). Western blots confirmed that nuclear levels of Stat3 were increased in WA314 infected compared to mock and WAC infected macrophages and were strongly decreased in WA314ΔYopM infected cells (Fig. 3C). Nuclear levels of Stat1 were elevated in all infection conditions (WAC, WA314 and WA314ΔYopM) compared to non-infected cells (S Fig. 3). Together these data suggest that YopM actively induces translocation of Stat3 to the nucleus. In order to test whether the YopM induced nuclear translocation of Stat3 contributes to the YopM triggered expression of *IL10* (Darnell et al. 1994), we knocked down Stat3 in the macrophages using siRNAs (siStat3(1); Dharmacon; 20nM and siStat3(2), Ambion; 20nM) (S Fig. 2B). siStat3(1) reduced WA314 induced IL-10 mRNA levels in macrophages from 3/3 donors when compared to a non-targeting control siRNA (Fig. 3D and S Fig. 2C). The inhibitory effect of siStat3(2) on WA314 induced IL-10 mRNA levels was detectable in 2/3 donors (Fig. 3D and S Fig. 2D).

**Figure 3.**
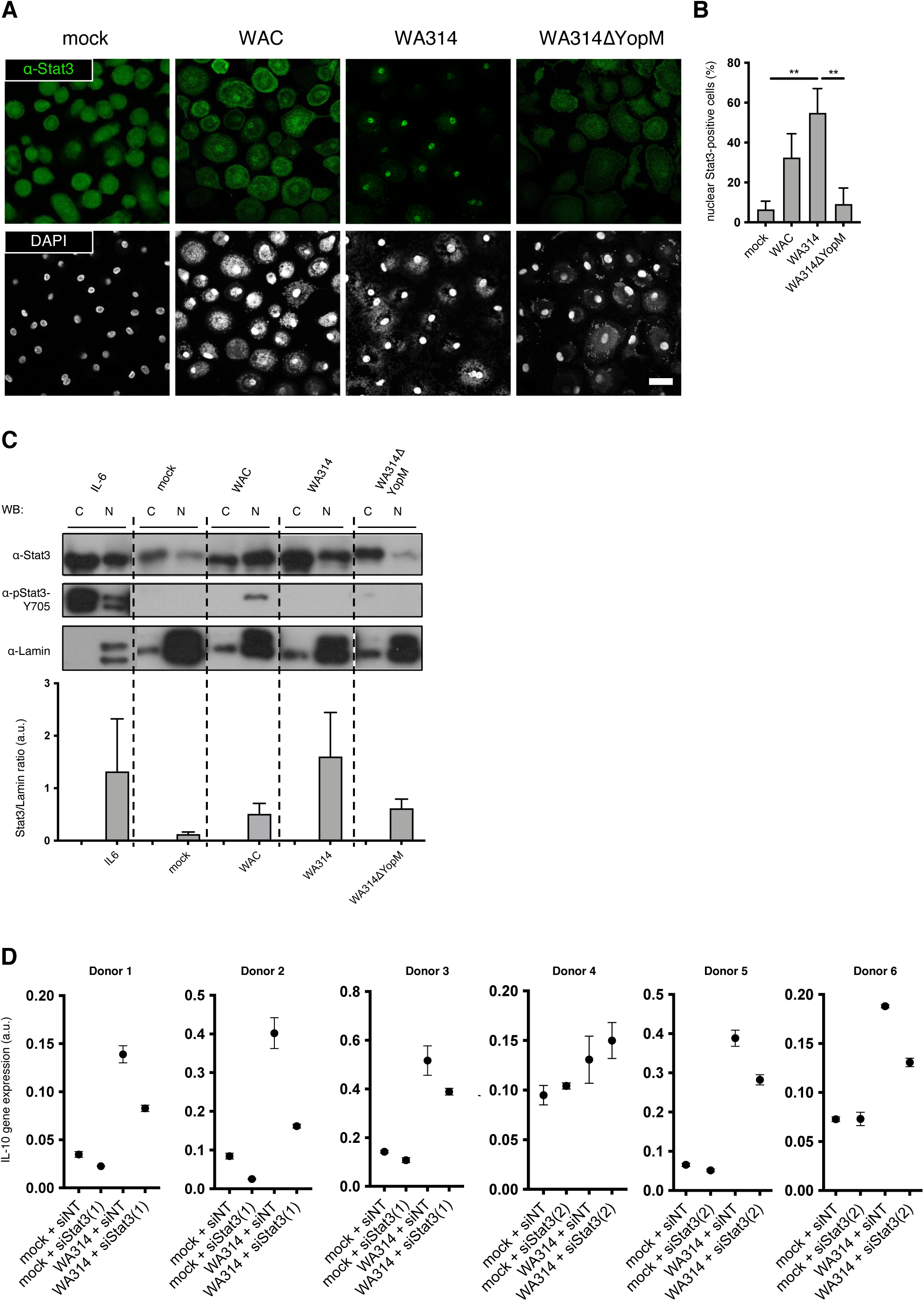
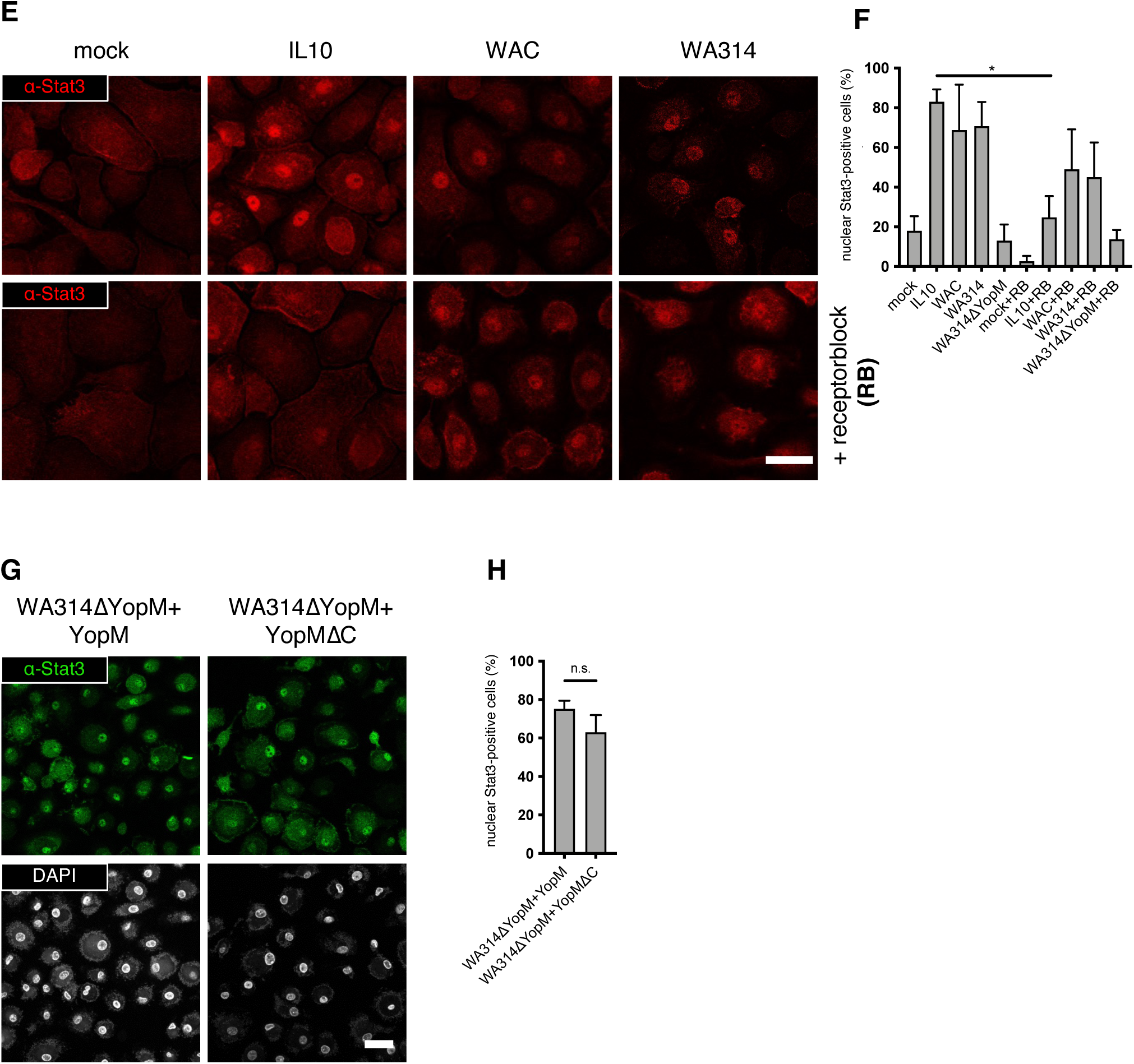
YopM induces nuclear translocation of Stat3. **A)** Human primary macrophages were infected with indicated *Yersinia* strains for 3h and immunofluorescence stained using anti-Stat3 antibody and DAPI **B)** Percentage of cells with nuclear Stat3 was determined. **p<0.01. **C)** YopM promotes Stat3 nuclear localization in the absence of phosphorylation on tyrosine residue 705. Macrophages were infected and cytosolic-(C) and nuclear (N) fractions were investigated by immunoblot using indicated antibodies. Band intensities of Stat3 signals were determined by ImageJ and intensity ratios of nuclear Stat3/Lamin were calculated. Each bar represents mean ± SEM of values from 6 experiments. **D)** siRNA mediated knockdown of Stat3 decreases IL-10 gene expression. Primary human macrophages were not treated (mock), treated with non-targeting (siNT), siStat3(1) Dharmacon) or siStat3(2) (Ambion) for 48 h and infected with WA314 for 5 h. Total RNA was subjected to quantitative RT-PCR. For each siRNA triplicate samples of macrophages derived from three different donors (Donor_1 to Donor_3; siStat3(1) and Donor_4 to Donor_6; siStat3(2)) were investigated, arbitrary units (a.u.). **E)** YopM promotes nuclear relocalization of Stat3 independently of autocrine secreted IL-10. Macrophages were left untreated (upper row) or incubated for 15min with human IL-10 receptor beta antibody (R&D systems; 25µg/ml) (bottom row), left uninfected (mock), stimulated with IL-10 (IL-10) or infected with indicated *Yersinia* strains. **F)** Quantification of nuclear Stat3-positive cells, *p<0.05. **G)** YopM promotes nuclear relocalization of Stat3 independently of its interaction with RSK. Cells were infected with Yersinia strains WA314ΔYopM+YopM and WA314ΔYopM+YopMΔC for 3h and immunostained with Stat3 antibody (green) or DAPI and **H)** quantified for nuclear Stat3 signal from three independent experiments with n=100 cells; **p<0.01.

We conclude that the YopM induced Stat3 translocation to the nucleus of human macrophages contributes to the enhanced expression of *IL10* stimulated by YopM.

To test whether autocrine IL-10 activating JAK-STAT signaling in the context of a positive feedback loop could be responsible for increased Stat3 nuclear translocation, we blocked the IL-10 receptor b (IL-10Rb) on the macrophage surface using an anti-IL-10Rb antibody (Fig. 3E; (O’Farrell et al. 1998)). As a positive control, we verified that anti-IL-10Rb blocked the IL 10 induced nuclear translocation of Stat3 (Fig. 3E and F). In comparison, WAC or WA314 induced nuclear translocation of Stat3 could not be inhibited by the anti-IL-10Rb antibody (Fig. 3E and F). This indicates that neither the WAC induced nor the WA314 induced nuclear translocation of Stat3 are mediated by an autocrine IL-10-triggered stimulation of JAK-STAT signaling.

JAKs phosphorylate Stat3 on Y705 causing its dimerization, nuclear translocation and DNA binding (Darnell et al. 1994, Ihle 1995). We asked whether the YopM induced increase in nuclear Stat3 may be due to increased Stat3 Y705 phosphorylation. As a positive control we verified that IL6, an established activator of JAK-STAT signaling, induced nuclear translocation and Y705 phosphorylation of Stat3 in the macrophages (Fig. 3C and S Fig. 2E and F). In WAC infected cells only a minor and in WA314 infected cells no Stat3 Y705 phosphorylation occurred concomitant with Stat3 nuclear translocation (Fig. 3C and S Fig. 2E and F).

To investigate whether the association of YopM with RSK (Berneking et al. 2016) is required for stimulation of nuclear translocation of Stat3, we infected macrophages with a YopM deletion strain recomplemented with i) C-terminally truncated YopM deficient for RSK binding (WA314ΔYopM+YopMΔC) or ii) wild-type YopM (WA314ΔYopM+YopM) (Berneking et al. 2016). Levels of nuclear Stat3 were similar in macrophages infected with strains expressing RSK binding deficient and wild type YopM (Fig. 3G and H). Furthermore, WA314 infected cells pretreated with the RSK kinase inhibitor LJI308 and WA314-infected untreated control cells displayed approximately the same level of nuclear Stat3 (S Fig. 2G). We conclude that YopM increases nuclear translocation of Stat3 independent of its interaction with RSK or activation of RSK.

In summary, our data suggest that YopM stimulates nuclear import of Stat3, which contributes to IL-10 gene expression. The exact underlying mechanism is unknown, but it is independent of preceding IL-10 production (excluding a positive feedback loop), Stat3 Y705 phosphorylation, and RSK activity. We propose that besides RSK activation, stimulation of Stat3 nuclear translocation constitutes another mechanism through which YopM increases expression of *IL10* in *Yersinia* infected macrophages.

## Discussion

Pathogenic *Yersinia* strains deficient in YopM show a dramatic reduction in virulence in mouse models, indicating that YopM is a critical pathogenicity factor (Trulzsch et al. 2004, McCoy et al. 2010, McPhee et al. 2010). The YopM protein contains a varying number of leucine-rich repeats depending on the *Yersinia* strain, through which it acts as a scaffold for multiple interaction partners, including kinases of the RSK and PKN/PRK families, the Rho GTPase effector IQGAP1, the DEAD box helicase DDX3 and caspase-1 (Leung et al. 1990, Skrzypek et al. 1996, Hines et al. 2001, Heusipp et al. 2006, LaRock et al. 2012). The interaction with RSK and PKN/PRK family kinases, leading to phosphorylation and activation of these kinases, has been shown to mediate distinct cellular effects of YopM. First, YopM prevents activation of the pyrin inflammasome by maintaining PKN activity in cases where Rho GTPases, the physiological activators of PKN, have been inactivated by other *Yersinia* effectors (Brodsky et al. 2010, Chung et al. 2016). Second, by binding to and activating RSK in the nucleus of host cells, YopM drives transcription of *IL10* (Berneking et al. 2016). Besides RSKs that act downstream of ERK 1/2 kinases (Cargnello et al. 2011), expression of the IL-10 gene can be induced by a multitude of other signaling pathways that connect to different transcription factors (Saraiva et al. 2010). Induction of IL-10 often occurs together with pro-inflammatory cytokines and may negatively regulate these pro-inflammatory cytokines (Saraiva et al. 2010). We found that IL-10 gene expression together with expression of dozens of proinflammatory cytokine genes is strongly induced by avirulent yersiniae in primary human macrophages. Notably, virulent bacteria, through its effector Yops, suppressed not only proinflammatory gene expression but also *IL10* expression almost completely, to a large extent through YopP. In this setting, YopM counteracts the inhibitory effects of YopP and the other Yops on IL-10 gene expression, and maintains an elevated level of IL-10 gene expression. Although this YopM effect appears small compared with the effect of, for example, LPS on IL-10 gene expression, IL-10 protein levels in macrophages are still twice as high after infection with YopM expressing than after infection with YopM-deficient bacteria. We propose that this elevated IL 10 level supports the bacterial infection process.

Here we wanted to get further insights into the mechanisms of YopM-induced IL-10 gene expression. For this we compared transcriptomic analyses of human macrophages infected with YopM-expressing and YopM-deficient *Yersinia* strains. We discovered that YopM causes significant changes in expression of JAK-STAT signal pathway genes, and that this is associated with nuclear translocation of the central transcription factor Stat3. Stat3 promotes the expression of immunosuppressive genes stimulated by the activated IL-10 receptor and also of the IL-10 gene itself as part of a positive feedback loop (Donnelly et al. 1999). These properties of Stat3 have probably made it an effective target of different pathogens that seek to suppress the immune response. Whereas *Toxoplasma gondii* activates Stat3 through its effector rhoptry kinase (Butcher et al. 2011, Duell et al. 2012), the Salmonella effector SarA/SteE triggers phosphorylation and activation of Stat3 through the cellular kinase GSK-3 (Gibbs et al. 2020). We found that also *Yersinia* YopM activates Stat3 which is reflected by its translocation into the nucleus, but that neither its known interaction with RSKs plays a role in this, nor does its stimulation of IL-10 production, which rules out a positive feedback loop in the JAK-STAT signaling pathway. Also, tyrosine phosphorylation of Stat3 was not altered by YopM. The question therefore arises by what mechanism YopM induces the nuclear translocation of Stat3. Because YopM itself translocates into the nucleus via an unknown mechanism, it could act as a carrier for unphosphorylated Stat3 (Skrzypek et al. 1998, Skrzypek et al. 2003, Scharnert et al. 2013, Berneking et al. 2016). YopM could also induce the production of a hitherto unidentified cytokine other than IL-10 that stimulates the JAK-STAT pathway in an autocrine fashion and thereby triggers Stat3 translocation.

Overall, this work is another example of a molecular mechanism that is hijacked by *Yersinia* YopM to rewire a cellular immune response to the advantage of the pathogen. Through this activity YopM maintains an elevated level of the immunosuppressive cytokine IL-10, which has been shown to be beneficial for a number of pathogens (Cyktor et al. 2011). It remains a task for future work to decipher the exact molecular mechanism of this unique YopM activity and how it contributes to the fascinating network of the immunosuppressive activities of YopM.

## Material and Methods

### Bacterial strains

*Yersinia enterocolitica* strains used in this study (S1 Table) are derivatives of the serotype O:8 strain WA314 harboring the virulence plasmid pYVO8. The virulence plasmid-cured derivative of WA314 was termed WA-C (Heesemann et al. 1983). WA314ΔYopM was constructed by replacing the YopM gene in WA314 with a kanamycin resistance cassette (Trulzsch et al. 2004). By complementation of WA314ΔYopM with isogenic full length YopM or C-terminal truncated YopM the WA314ΔYopM+YopM and strain WA314ΔYopM+YopMΔC were created (Hentschke et al. 2010). WA314ΔYopP was constructed by replacing the YopP gene with a chloramphenicol resistance cassette (Trulzsch et al. 2004).

### Cell culture

Human peripheral blood monocytes were isolated from buffy coats, kindly provided by Frank Bentzien (University Medical Center Eppendorf, Hamburg, Germany), as described previously (Kopp et al. 2006). The Ethical Committee of the Ärztekammer Hamburg, Germany approved the analysis of anonymous blood donations (WF-015/12). Cells were used for *Yersinia* infection after one-week culture in RPMI1640 containing 20% autologous serum.

For infection of the primary human macrophages, culture medium was changed to RPMI1640 16h before infection. *Yersinia* cultures were grown overnight at 27 °C, diluted 1:20 in fresh Luria-Bertani (LB) broth and grown for another hour at 37 °C. Bacteria were resuspended in ice-cold PBS containing 1mM MgCl_2_ and CaCl_2_ and added to at a multiplicity-of-infection (MOI) of 100 for indicated time points.

### RNAi

Human Stat3 (Gene ID: 6774) siRNA smartpool M003544 (siRNA-IDs: D-003544-02, target sequence GGAGAAGCAUCGUGAGUGA, D-003544-03, target sequence CCACUUUGGUGUUUCAUAA, D-003544-04, target sequence UCAGGUUGCUGGUCAAAUU, D-003544-19, target sequence CGUUAUAUAGGAACCGUAA), Ambion silencer select IDs744 (siRNA-IDs: D-003544-02, target sequence GGCUGGACAAUAUCAUUGAtt) and control Non-Targeting siRNA (Pool #2 targeting firefly luciferase and EGFR) were from the Dharmacon siRNA collection (Lafayette, USA). siRNA transfection of human primary macrophages was performed using the Neon Transfection System (Invitrogen, Darmstadt, Germany) with standard settings (1000 V, 40 ms, 2 pulses) and 1.5 μg of siRNA 48 h before infection of the cells.

### Library preparation and high-throughput sequencing

Total RNA of human macrophages was isolated according to manufacturer’s instructions using the RNeasy extraction kit (Qiagen). mRNA was extracted using the Next Poly(A) mRNA Magnetic Isolation module (New England Biolabs; NEB) and RNA-seq libraries were generated using the Next Ultra RNA Library Prep Kit for Illumina (NEB) as described before (Berneking et al. 2016).

Libraries were assessed for size and quality using a BioAnalyzer High Sensitivity Chip (Agilent) and sequenced on the Illumina HiSeq 2500 instrument with 39.1-55.8 million reads per sample (single read 51 bp run).

Alignment of the reads to the human reference assembly (GRCh38.97) was performed using STAR (Dobin et al. 2013). To obtain the number of reads mapping to each gene FeatureCounts (Huang et al. 2014) was employed. Statistical analysis of differential expression was carried out with DESeq2 (Love et al. 2014). Raw counts were normalized and variance-stabilized using the regularized log transformation (rlog) from the DESeq2 package (Love et al. 2014). To remove the batch effects for plotting purposes, we extracted normalized and variance-stabilized expression values from DESeq2 and corrected them for batch effects using removeBatchEffect from the limma package (Ritchie et al. 2015) to allow for sample comparison with principal component analysis (PCA) and clustered heatmaps. In the end, DESeq2 was applied again to identify differentially expressed genes (DEGs) between conditions. Comparison to one public dataset (GSE43700; (Teles et al. 2013)) revealed that approximately 5% of DEGs in WAC, WA314 and WA314ΔYopM infected cells (FDR < 0.05, > 1-fold change; 189/2972; 108/1800, 40/926, respectively) compared to non-infected cells correspond to IL-10-dependent genes RNA-seq data have been deposited in the ArrayExpress database at EMBL-EBI (www.ebi.ac.uk/arrayexpress) under accession number E-MTAB-10473. RNA-seq data from additional replicates of mock, WA314 6h and WA314ΔYopM 6h were obtained from European Nucleotide Archive (ENA) at http://www.ebi.ac.uk/ena/data/view/PRJEB10086.

### Clustering and correlation analysis

Clustering analysis of all IL-10-dependent DEGs from comparisons was performed with rlog counts in R (software 3.2.3 5; R Core Team (2017), R Foundation for Statistical Computing, Vienna, Austria, https://www.R-project.org/.) with pheatmap package. Clustering was performed with clustering distance based on Pearson correlation and Complete clustering method. Clustering distance, clustering method and number of clusters were selected so that all meaningful clusters were identified by the analysis. For heatmaps rlog counts from DESeq2 analysis were scaled by row (row Z-score) indicated by blue-white-red color gradient. All replicates were used for the analysis but just 2 representative replicates were shown for each sample. KEGG pathway analysis was performed with Webgestalt (http://www.webgestalt.org) and heat map of KEGG pathways (Fig. 2C) was generated with Morpheus (https://software.broadinstitute.org/morpheus/).

### TF motif analysis

TF motif enrichment for known motifs was performed using HOMER (http://homer.ucsd.edu/homer/motif/ (Heinz et al. 2010)). Command findMotifs.pl was used and a list of gene symbols was supplied as an input. Motifs were searched in the region 400 bp upstream and 100 bp downstream of the TSS by specifying parameters -start -400 -end 100. For presentation of enriched TF motifs results from known motifs were used.

### Quantitative real time PCR (RT-qPCR)

Uninfected or infected primary human macrophages were subjected to RNA extraction using the RNeasy extraction kit (Qiagen). Two µg of RNA each was reverse transcribed with the iScript cDNA Synthesis Kit (Bio-Rad). RT-qPCR reaction were performed using the TaqMan® Fast Advanced Master mix (Applied Biosystems). IL-10 mRNA levels were determinated using gene specific primers (Hs00961622_m1; ThermoFisher Scientific). As reference the expression of housekeeper genes GAPDH (Hs02758991_g1), TBP (Hs00427620_m1) and B2M (Hs00187842_m1) were analyzed (ThermoFisher Scientific). RT-qPCR was performed with the LightCycler^®^ 480 Instrument (Roche Life Science) and data were analyzed according to manufacturer’s instruction (Roche LightCycler® 480 software; release 1.5.1.62). Reference genes and external standards were employed for the relative quantification of different expressions. Statistical analysis was performed with a linear mixed model taking into account random intercept. Data was transformed to ln-Score to ensure normal distribution using IBM® SPSS®-Software.

### Western blotting of macrophage cell lysates

Primary human macrophages were washed once with ice cold PBS prior to harvest of cells with TBS supplemented with protease inhibitors (Complete®, Roche) and phosphatase inhibitors (PhosStop®, Roche). Total protein levels were quantified by Bradford protein assay kit. Equal amounts of protein were separated by SDS-PAGE and transferred to polyvinylidene difluoride (PVDF) membrane (Immobilion-P, Millipore, Schwalbach, Germany) by semi-dry blotting. The membrane was incubated with blocking solution (5% milk powder (w/v) or 3% Bovine serum albumin (BSA) (w/v) in TBS supplemented with 0.01% Tween 20; TBS-T) for 30 min and subsequently with primary antibodies at 4°C overnight. Primary antibodies used in this study were monoclonal anti-RSK1 (C-21) antibody (Santa Cruz Biotechnology), mouse anti STAT3 antibody Stat3 (Cell signaling), rabbit anti-STAT3 antibody Stat3 (Cell signaling), rabbit anti-phospho-STAT3 (Tyr705) antibody Stat3 (Cell signaling), rabbit anti-STAT1 antibody Stat1 (Cell signaling), monoclonal rabbit Lamin A/C (Cell Signaling), monoclonal mouse anti-Histon3 (H3, Cell signaling), monoclonal rabbit anti-phospho-H3S10 (Abcam) (dilution 1:1000); anti-phospho-S380RSK (Cell Signaling) (dilution 1:2000); polyclonal anti GAPDH (Sigma-Aldrich), monoclonal anti-actin (Millipore, Schwalbach, Germany); (dilution 1:2500). Secondary antibodies were horseradish peroxidase linked sheep anti-mouse IgG (GE Healthcare) and donkey anti-rabbit IgG (Cell signaling, GE Healthcare) used in a dilution of 1:10000 and incubated at room temperature for 2 h. Washing steps were performed with TBS-T to minimize unspecific binding. Antibody signals were visualized with chemiluminescence technology (Supersignal West Femto, Pierce Chemical, Rockford, USA) and captured on X-ray films (Fujifilm, Düsseldorf, Germany). Developed films were scanned (CanonScan 4400F (Canon, Tokio, Japan) to quantify protein band intensity. Quantification of signal intensity of scanned films CanonScan 4400F (Canon, Tokio, Japan) was performed using ImageJ analysis software Version 1.43u (National Institute of Health, NIH).

### Immunofluorescence staining and confocal microscopy

Human peripheral blood monocytes-derived macrophages grown on glass coverslips (Marienfeld GmbH, Lauda-Königshafen, Germany) were fixed with 3.7% (v/v) formaldehyde in PBS for 5 min, permeabilized with Methanol for 6 min, blocked with 5% BSA in PBS for at least 15 min and incubated with a 1:100 dilution of primary antibody for 1 h followed by a 1:200 dilution of secondary antibody for 45 min. After three wash steps in PBS, coverslips were mounted in Mowiol (Calbiochem, Darmstadt, Germany). Images were acquired with a confocal laser scanning microscope (Leica DMI 6000 with a Leica TCS SP5 AOBS confocal point scanner) equipped with a 63 x oil immersion HCX PL APO CS objective (NA 1.4–0.6). Acquisition was performed with Leica LAS AF software (Leica Microsystems, Wetzlar, Germany). Volocity 6 software (Improvision, Coventry, UK) was used for processing of images.

### ELISA

We measured levels of secreted IL-10 in supernatants from infected or non-infected human primary macrophages cells using commercially available enzyme-linked immunosorbent assay (ELISA) kit (R&D Systems) according to the manufacturers’ instructions.

### Statistical analysis

Statistical analysis was performed using Graph pad prism version 6.0. At least three independent experiments were compared by paired t-test, one-way ANOVA with Bonferronís post-test or two-way ANOVA if not indicated otherwise. p-values < 0.05 were considered statistically significant.

## Suppl. figures

**Suppl. Figure 1.**
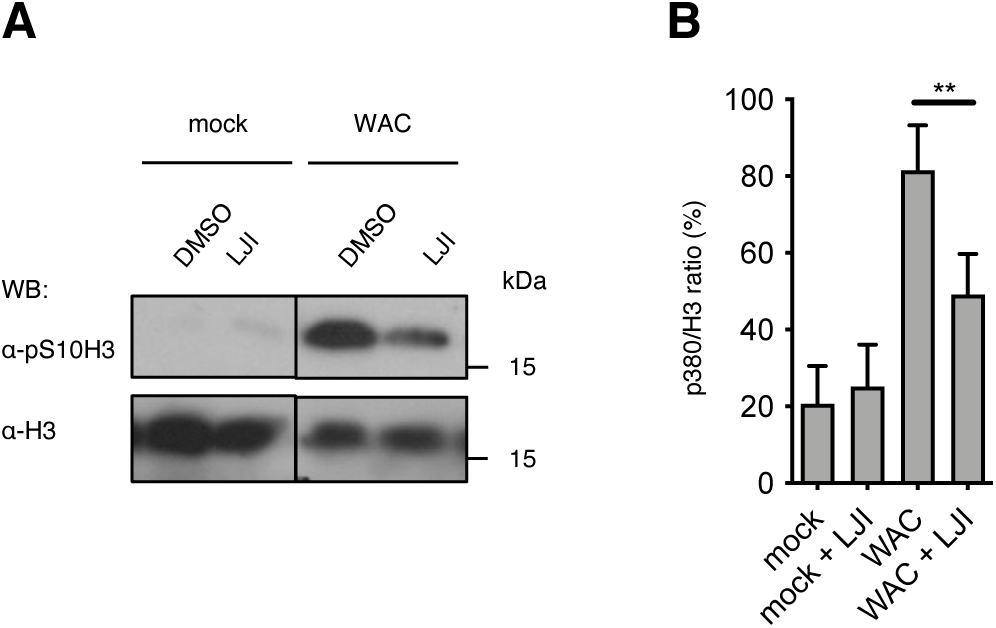
Inhibition of RSK abolishes phosphorylation of histone 3 at serine 10. **A)** Primary human macrophages were treated with DMSO (Sigma) or RSK inhibitor LJI (10µM, Sigma), infected with *Yersinia* WAC for 5h and subjected to Westernblot analysis. **B)** Phosphorylation of Histone 3 at Serine 10 residue was evaluated by quantification of band intensities to determine ratio of pS10/H3.

**Suppl. Figure 2.**
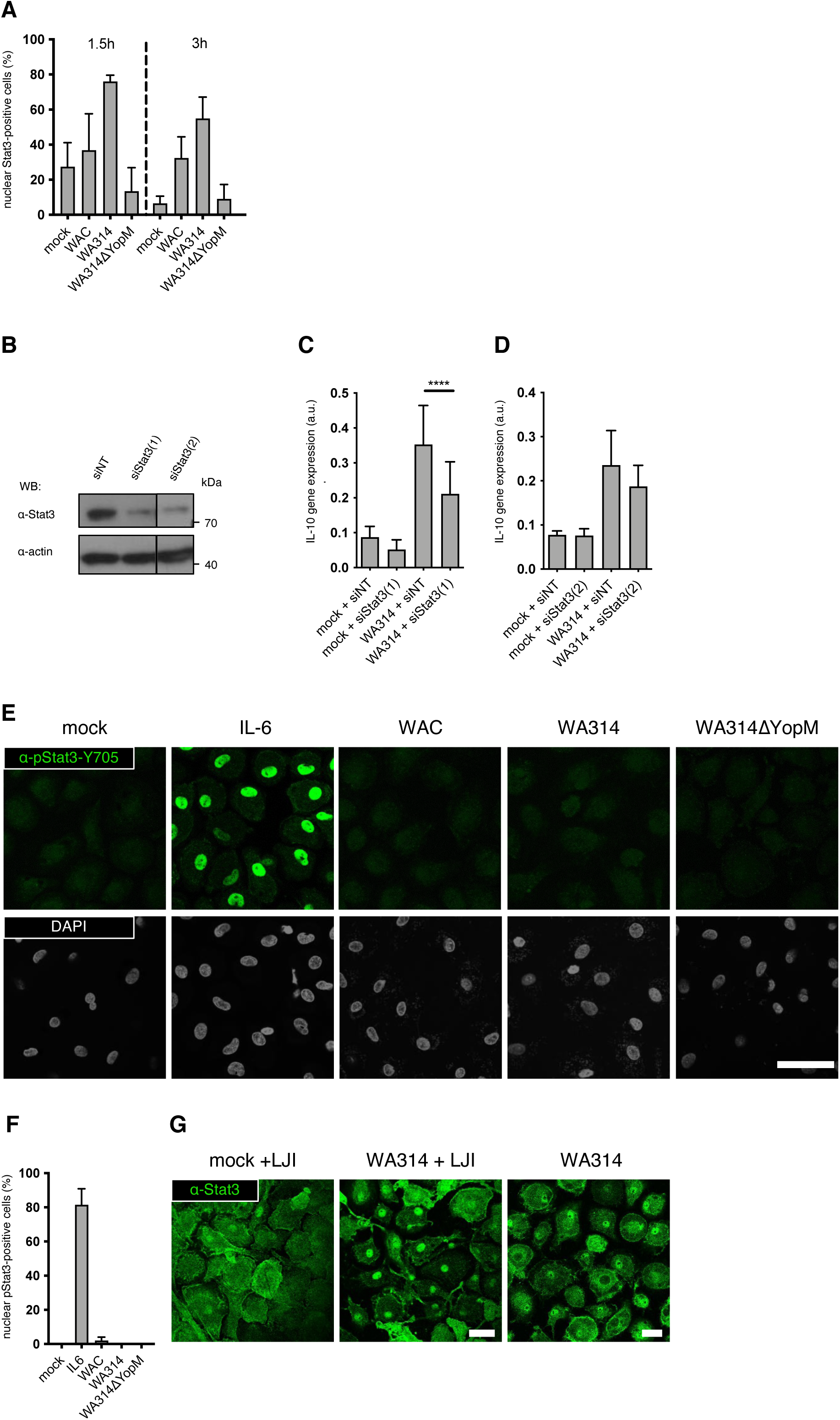
*Yersinia* activates Jak Stat3 signaling. **A)** Quantification of cells with nuclear Stat3 signal from three independent experiments with n=100 cells infected for 1.5 and 3h. **B)** Effective knockdown of Stat3 in different donors. Primary human macrophages were treated with siNT or siStat3(1) (Dharmacon; 20nM) and siStat3(2) (Ambion; 20nM) and **C)** infected with *Yersinia* wild type WA314 for 5h after 48h or left untreated (mock). Total RNA was isolated and analyzed by rtPCR for expression of *IL10*. **D)** Expression of *IL10* in cells treated with siNT or siStat3(2) that were mock or WA314 infected. Each Bar in graph represents mean ± SEM of values from 3 different donors; ****p<0.0001, arbitrary units (a.u.). **E)** Cells were infected with *Yersinia* strains WAC, WA314, WA314ΔYopM for 3h, stimulated with IL6 for 30 min or left untreated (mock), permeabilized and immunostained with phospho-Stat3 antibody (green) or DAPI. **F)** Quantification of cells with phosphorylated nuclear Stat3 signal. **G)** Cells were treated with RSK inhibitor LJI (10µM, Sigma) for 1h prior to infection with Yersinia WA314, WA314ΔYopM or left mock-infected and immunostained with Stat3 antibody (green) or DAPI. Scale bar, 20µm.

**Suppl. Figure 3.**
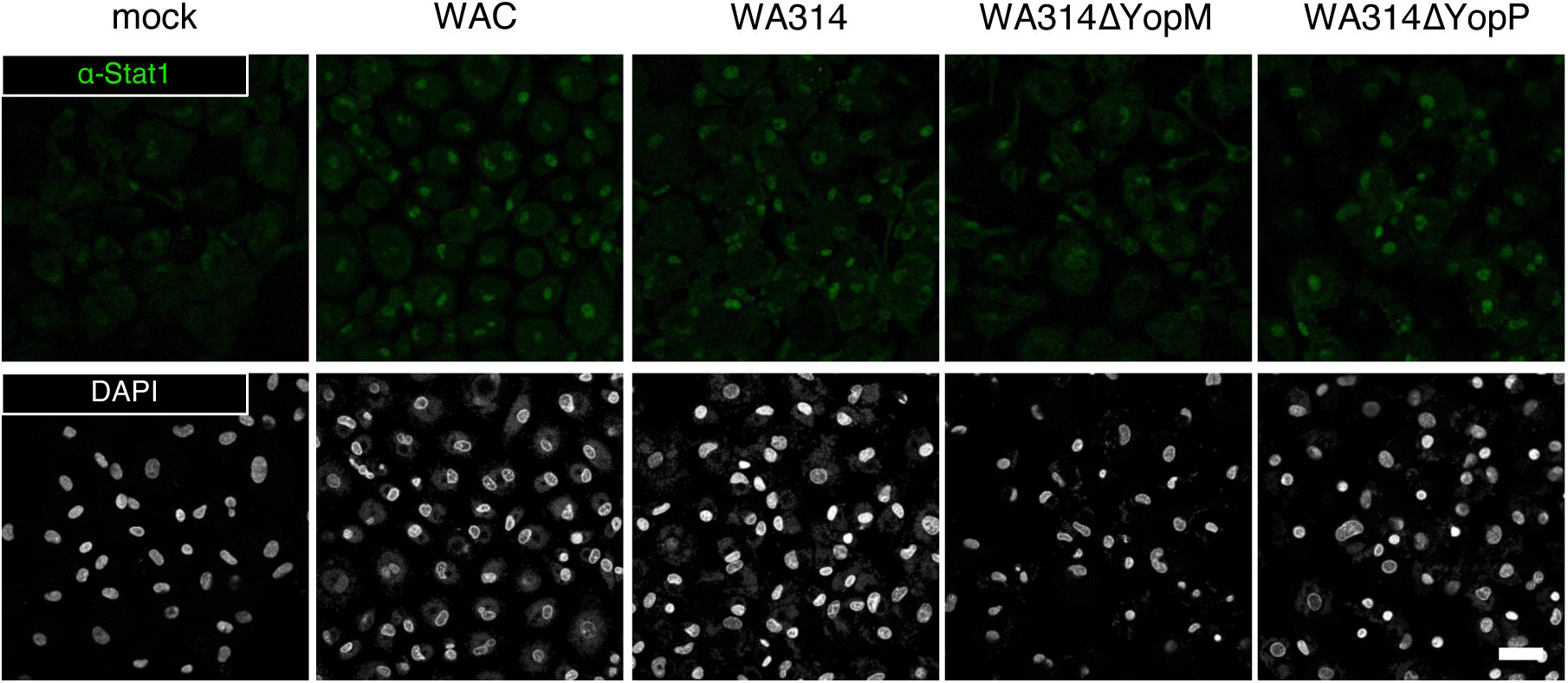
*Yersinia* activates Jak Stat1 signaling. Macrophages were uninfected (mock) or infected with *Yersinia* strains WAC, WA314, WA314ΔYopP and WA314ΔYopM for 3h and immunostained with Stat1 antibody (green) or DAPI.

